# Intentions of Canadian health professionals towards recommending exercise to people living with ALS

**DOI:** 10.1101/648790

**Authors:** Kelvin E Jones, Tanya R Berry, Vanina Dal Bello-Haas

## Abstract

**Background:** To provide a nationwide overview of the attitudes, social pressure, perceived ability and intentions of health professionals toward exercise prescription for people living with ALS (pALS).

**Methods:** An online survey of physician and other health professionals (HPs) working in academic ALS clinics across Canada.

**Results:** The response rate was 48% (84/176) with 30% of respondents identifying as physicians, 63% as other HPs and the remainder as administrative or research personnel. Respondents were sharply divided in their intentions to provide exercise counsel: 24% unlikely and 45% likely. Respondents with low intentions were HPs that considered this activity outside their scope of practice. Measures of intention and attitude were more positive for flexibility compared to strength and aerobic exercise. Perceptions of social pressure and ability to provide exercise counsel were significantly correlated with intention across the three exercise modes in all respondents. Qualitative themes identified as barriers to exercise prescription were lack of confidence or competence (31% physicians, 32% HP), patient tolerance (30% HP), lack of evidence (22% physicians) and lack of infrastructure (22% physicians).

**Conclusions:** While “lack of evidence” for the benefit of exercise was a deterrent for physicians, the larger issue for all respondents was building competence and confidence in exercise prescription for pALS.

## Background

The American College of Sports Medicine (ACSM) manages a global health initiative called Exercise is Medicine^®^ (EIM). The central belief of EIM is that physical activity and exercise prescription are an integral component of multidisciplinary healthcare. EIM advocates for the benefits of exercise for individuals living with chronic disease or disability including those with neurologic and neuromuscular disorders. Evidence-based guidelines for exercise management in people living with ALS (pALS) are available as part of this initiative [1] but it is not clear what the current beliefs about exercise prescription are for pALS or whether health professionals are aware of these guidelines. Traditionally, sport- and work-related exercise was considered a potential risk factor for sporadic ALS [2, 3, 4, 5, 6, 7]. At the time the ACSM guidelines were published, a review of the epidemiological litera-ture concluded that evidence of risk from exercise “is limited, conflicting and not of sufficient calibre to allow undisputed conclusions” [8].

The most recent epidemiological evidence concludes that recreational physical activity may be protective rather than a risk factor [9, 10, 11] but a possible direct association between strenuous exercise and risk for ALS mortality remains [12, 13]. The most recent Cochrane review on the topic points out the need for more rigorous randomized controlled trials (RCT) while acknowledging the moderate benefit of exercise and lack of reported adverse effects in existing RCTs [14]. Since that review, but prior to our survey, two trials have been completed that found a moderate intensity (60% of maximal power output) exercise program reduced the rate of motor deterioration in pALS [15] and that resistance and endurance exercise are safe for pALS [16]. While the evidence for modest beneficial effects of exercise on the progression of ALS accumulates, the lack of reported adverse effects in RCTs is consistent. However, anecdotal experiences conveyed by pALS on the PatientsLikeMe website [17] as well as published editorial commentary [18] suggest that this issue remains controversial and that there may be a lingering perception that exercise is contraindicated for pALS.

Given the current state of evidence about physical activity and disease progression in ALS, we sought to determine the opinions of front-line ALS health professionals on this issue. Since health professionals play a vital role in motivating change in the physical activity of patients, their intentions are an important measure to compare with the increasing call for exercise counsel in the clinic [19, 20]. The primary goal of the survey was to measure their intention to provide exercise counsel and the factors that contribute to that decision. The secondary objectives were to (i) determine whether mode of exercise (i.e. flexibility, strength, aerobic) influenced intentions and (ii) gauge differences between physicians and other health professional respondents. Participants were asked to consider the ACSM guidelines for pALS in their responses which are targeted to individuals in the early stages of ALS when functional impairments are moderate. It was proposed that the intention to provide exercise counsel would be low to moderate, in accord with anecdotal reports of pALS and health professionals, and that intentions would be associated with the constructs of the theory of planned behavior (TPB), as described in the methods section [21]. It was also proposed that intentions and attitudes towards flexibility-type exercise would be more positive compared to strength and aerobic exercise modalities since stretching is a common practise in ALS management [22].

## Methods

The procedures were approved by the University of Alberta’s Research Ethics Board #2 (Pro00047048) and were in accordance with the ethical standards outlined in the “Tri-Council Policy Statement” and the Helsinki Declaration of 1975, as revised in 1983. Potential respondents were informed that survey completion was considered implied consent prior to participation.

### Participants

Administrative personnel associated with all 15 academic and 2 satellite ALS clinics across Canada were contacted to develop a sampling frame. The number of staff and their employment categories were collected to estimate the total number of potential respondents as 176. Of the total number of potential respondents, 48 were classified as physicians and 104 as non-physician health professionals (HPs) with the remaining 24 individuals classified as administrative, support or research staff. The sampling frame was considered a current list of Canadian health professionals working directly with people living with ALS in Spring 2015. Participants were recruited to complete an anonymous online survey by distribution of an advertisement through the Canadian ALS Research Network and through internal email via the provincial ALS clinic administrators. Hard copy advertisements were sent to the clinics along with reminder emails on a weekly basis. Recruitment took place over a 4-week period, January-February 2015. The open survey link, hosted by SurveyGizmo (www.surveygizmo.com), was an active link on all electronic communication and advertisements disseminated to team members. A copy of the survey and the advertisememtn is available online at Figshare https://doi.org/10.6084/m9.figshare.8044622.v1.

### Instrument development

The survey tool was developed using the theory of planned behavior (TPB) [21]. The TPB states that intentions precede behavior and these intentions are influenced by three constructs: attitude (an individual’s positive or negative judgment of the behavior), subjective norms (an individual’s perception about whether their peers think they should do the behavior and if others like them do the behavior), and perceived behavioral control (the individual’s evaluation of their confidence to perform the behavior in conjunction with their control over doing the behavior). In this case the behavior in question is ‘exercise counsel’ or ‘exercise prescription’. The TPB is suitable to examine psychological constructs that might influence a health provider’s intentions to use clinical guidelines with patients [23]. Previous research has shown that perceived behavioral control was the strongest predictor of doctor’s intentions to use clinical practice guidelines whereas subjective norms was the most important for nurses and other health professionals [23]. A questionnaire was developed consisting of multi-item open and closed questions. Pilot testing was done to determine content validity, readability and to assess the criterion validity of the constructs in the survey [24]. The pilot test sample included nine health professionals with knowledge of neuromuscular disorders, but not connected to the target ALS clinics: three physicians (physiatry, neurology) and six other HPs (speech language pathologist(s), nurse(s), physical and occupational therapist(s)). Participants in the pilot test completed the web survey while simultaneously verbalizing thoughts using the “think aloud technique” [25]. Both quantitative and qualitative data collected during the pilot test were used to refine the survey instrument.

### Survey measures

Respondents answered seven questions to characterize demographics and their work environment. These included two validated questions to asses the level of work and leisure physical activity that were converted into a physical activity level value [26].

The survey items addressing the constructs of the theory of planned behavior asked for separate responses to three exercise modes: strength training, aerobic training and flexibility. The participants were asked to use the following ACSM guidelines for exercise in people living with ALS as a reference when responding to these survey items.

*”These guidelines stress individualized programming and are intended for early stage ALS. The ACSM guidelines recommend aerobic exercise 3 times a week for up to 30 minutes at an intensity of 50-80% of their age predicted peak heart rate. Strength training should be done on non-aerobic days at a low to moderate intensity with a load that allows 8-12 repetitions for 1-2 sets in good form. Flexibility exercises are recommended to be performed 1-2 times everyday.”* [1]

Responses were scored using Likert scales ranging from 1 to 7 with an option to respond, “Do not know” for each item. Anchors for the Likert scales included: very useless – very valuable, strongly disagree – strongly agree, very false – very true, and very unimportant – very important. Some item statements were negatively worded and were reverse coded prior to analysis.

#### Intentions

Intentions were assessed with the question: “*I will provide exercise counsel to my patients with ALS in the future*”, rated on a 1 (very unlikely) to 7 (very likely) scale. The score categorized the valence of response: positive (> 5.0), neutral (3.0 – 5.0), and negative (< 3.0). This descriptive categorization was also used for the following TPB constructs.

#### Attitudes

Attitude toward exercise for people living with ALS was assessed by response to four statements (e.g. *Exercise for people living with ALS is*: rated from 1 very useless to 7 very valuable). Internal consistency was good for strength (Cronbach’s *α* = 0.82) and aerobic (*α* = 0.71) training; therefore a mean score of these items was used to calculate a scale for attitude. Internal consistency for flexibility was lower (*α* = 0.63). We determined that the low value was due to the narrow range of responses on these items and a ceiling effect (i.e. 92% of responses were either a 6 or 7 on the Likert scale), so a mean score was used to calculate a scale for flexibility attitude as for the other two exercise modes. Higher values on the scale (maximum of 7) represent a more positive attitude.

#### Subjective Norms

The perception of peer beliefs toward exercise for people living with ALS was assessed by the response to four items (e.g. *My colleagues think exercise for people living with ALS is*: rated from 1 very unimportant to 7 very important). Internal consistency was good (strength *α* = 0.81, aerobic *α* = 0.70) to fair (flexibility *α* = 0.67); therefore a mean score of the items was used to create a scale for subjective norms. Higher values on the scale (maximum 7) represent a perception of positive peer belief.

#### Perceived Behavioral Control

Four items (Table S1) were used to assess an individual’s evaluation of their ability to provide exercise counsel. Internal consistency ranged from *α* = 0.59 (aerobic) to 0.62 (flexibility). Cronbach’s alpha was recalculated with each item deleted, but this procedure did not identify a specific problem item. Given the low internal consistency (< 0.7 [27]), single item as well as a mean score analysis was done. Higher values on the PBC scale indicate that an individual perceives a greater ability to provide exercise counsel.

#### Non-TPB Measures

There were seven additional items that evaluated participants’ response to the primary research question. Participants were asked whether exercise counsel was within their scope of practice. If they answered ‘yes’, a contingent item was presented that asked, “How much clinical team involvement is used when prescribing exercise”, and answered from 1 (completely collaborative) to 7 (entirely autonomous) for each of the three exercise modalities. Participants were asked if they were familiar with the ACSM exercise guidelines, whether they used the guidelines, what proportion of patients asked about exercise, and what proportion are capable of exercising according to guidelines. Participants were then asked how many patients they provided exercise advice to over the last month to provide a measure of prior behavior. The final item was an open-ended question that asked for up to 3 reasons why exercise counsel was **not** provided to all of their patients with ALS. A content analysis was done and five categories of responses were established iteratively with input from all authors. Two reviewers independently classified a randomized list of responses from participants and the inter-rater reliability was calculated (Cohen’s kappa = 0.91).

### Statistics

Data were analyzed using IBM SPSS version 23 software. Proportions and medians were calculated for nominal and ordinal variables. Continuous variables were described with mean and standard deviations, as well as tested for normality (Shapiro-Wilk test) prior to any parametric statistical tests. Alternate non-parametric tests were chosen if normality assumptions were violated. Spearman’s correlations were used for ordinal variables and Pearson’ product moment correlations for continuous variables after checking assumptions. Null hypothesis significance testing for differences in proportions was done using chi-square (χ^2^) test of homogeneity with post-hoc z-test. A Friedman two-way analysis of variance by ranks, for repeated measures, was used to test for differences between intentions to provide exercise counsel for strength, aerobic and flexibility exercise followed by pairwise comparisons using Wilcoxon signed-rank test. Other tests for group differences included Kruskal-Wallis H test (e.g. ACSM Familiarity across 3 independent groups) with post-hoc pairwise comparisons using Dunn’s procedure [28], two-way mixed ANOVA (e.g. attitude across three exercise modalities, between groups of respondents) and Student t test (e.g. years working in ALS field). Bonferroni correction was used to correct for multiple post-hoc comparisons and adjusted p-values reported. Multiple regression analysis was done to predict intentions from profession, and the three TPB constructs: attitude, subjective norms and perceived behavioral control. Linearity, homoscedasticity, and normality were assessed visually using partial regression plots, studentized residuals against predicted values and Q-Q plots. Independence of residuals was assessed by the Durbin-Watson statistic. Significance is reported using two-sided tests if p values are ≤ 0.05(*) or ≤ 0.01 (**).

## Results

Responses from 84 health professionals from all of the Canadian academic ALS clinics were received, representing an estimated response rate of 48% (sampling frame of 176). The respondents were divided into physicians (n=25) and nonphysician health professionals (n=53). The data from six respondents, categorized as administrative or research staff, were not included in further analysis due to low numbers. The demographics of the sample are summarized in Table 1. While there was no difference in the number of years employed in their profession, the physician group had significantly more experience working specifically with pALS: 14.3 ± 11.1 compared to 7.8 ± 7.4 years for other HP. A greater proportion of physicians reported that exercise counsel was within their scope of practice (88%, n=22) compared to other HP (44.2%, n=23, chi-square test of homogeneity, p < 0.0005). Other HP were divided into two groups based on this question for further analysis: *HP_Yes_* and *HP_No_*.

**Table 1.**
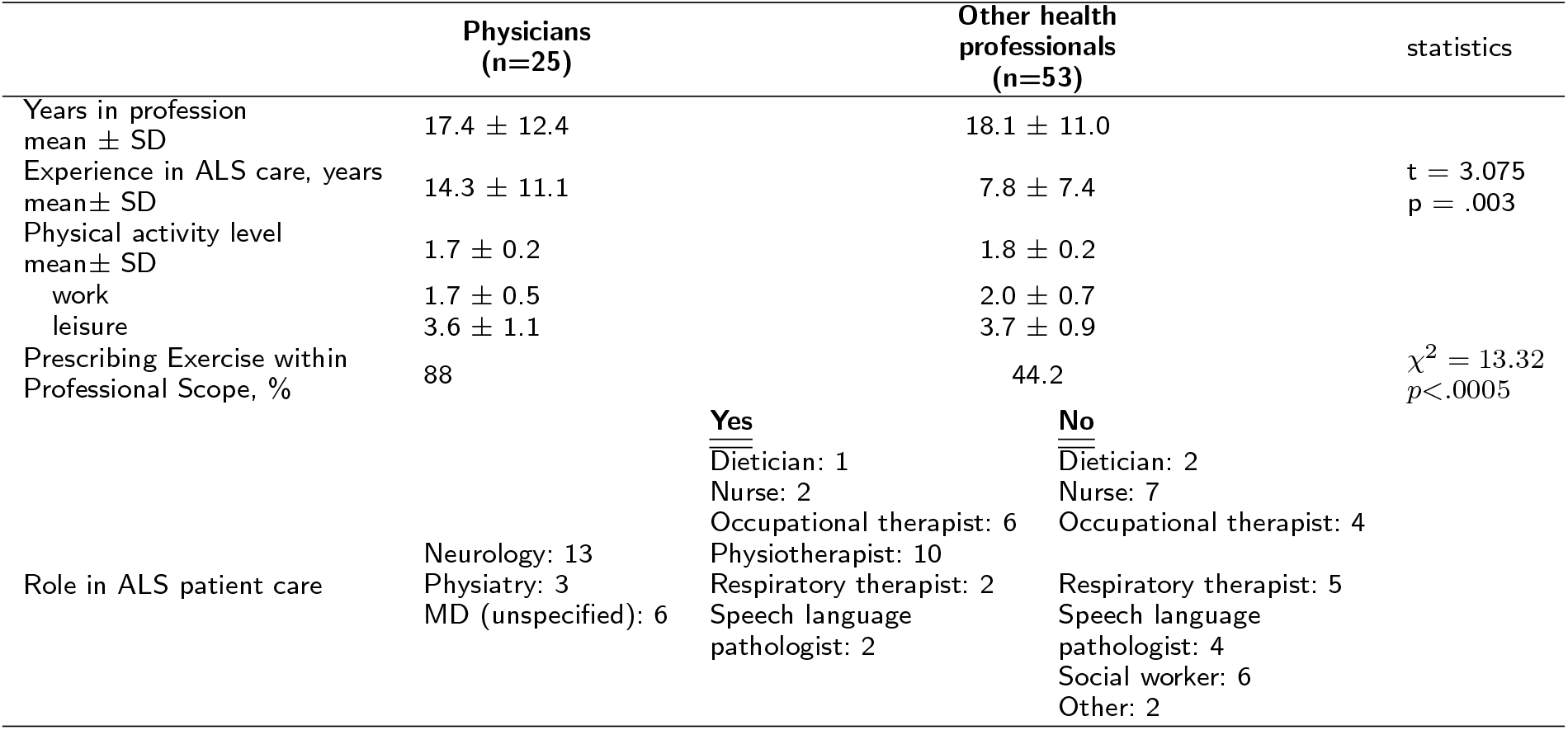
Demographics of Respondents

|  | Physicians<br>(n=25) | Other health<br>professionals<br>(n=53) | statistics |
| --- | --- | --- | --- |
| Years in profession<br>mean ± SD | 17.4 ± 12.4 | 18.1 ± 11.0 | |
| Experience in ALS care, years<br>mean ± SD | 14.3 ± 11.1 | 7.8 ± 7.4 | t = 3.075<br>p = .003 |
| Physical activity level<br>mean ± SD | 1.7 ± 0.2 | 1.8 ± 0.2 | |
| work | 1.7 ± 0.5 | 2.0 ± 0.7 | |
| leisure | 3.6 ± 1.1 | 3.7 ± 0.9 | |
| Prescribing Exercise within<br>Professional Scope, % | 88 | 44.2 | $\chi^2 = 13.32$<br>p < .0005 |
| Role in ALS patient care | Neurology: 13<br>Physiatry: 3<br>MD (unspecified): 6 | Yes<br>Dietician: 1<br>Nurse: 2<br>Occupational therapist: 6<br>Physiotherapist: 10<br>Respiratory therapist: 2<br>Speech language<br>pathologist: 2<br>No<br>Dietician: 2<br>Nurse: 7<br>Occupational therapist: 4<br>Respiratory therapist: 5<br>Speech language<br>pathologist: 4<br>Social worker: 6<br>Other: 2 |  |

### Topic of exercise and ACSM guidelines

A high proportion of pALS ask their healthcare professionals about exercise, but they are selective about who they discuss this topic with (Figure 1A). A majority of physicians (55%) and *HP_Yes_* (64%) report that exercise is discussed by more than 61% of pALS. In contrast, the majority of *HP_No_* (73%) report that exercise is discussed with 20% or fewer pALS. Few health professionals reported moderate-to-strong familiarity with the ACSM guidelines for exercise with pALS (Figure 1B). The median value for familiarity was significantly lower for the *HP_No_* group compared to the other two groups, which were not significantly different from each other (Kruskal-Wallis H test and post-hoc analysis details in Figure 1 legend). When asked whether participants used the ACSM guidelines during consultations with pALS the median response was 1.0 “Never” and there were no differences across the three groups (data not shown).

**Figure 1.**
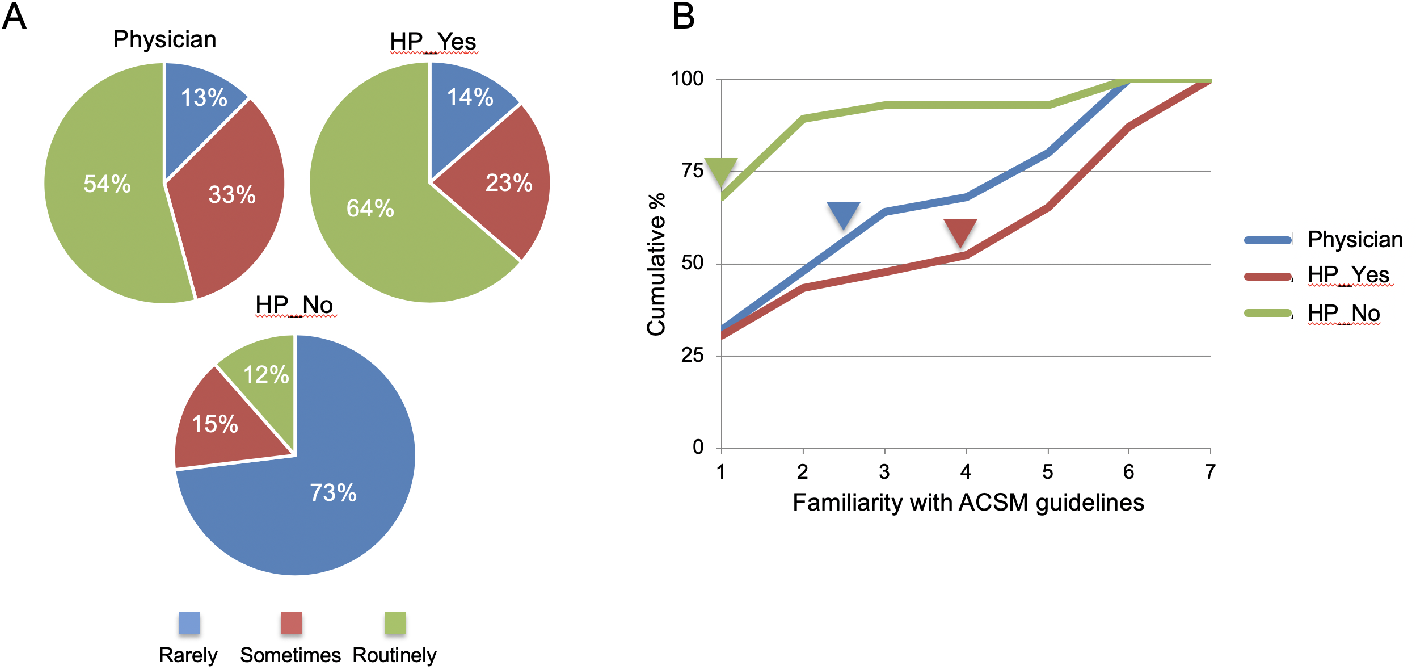
The question of exercise and guidelines. (A) Proportion of pALS that ask their healthcare professional about exercise. Categories of response: Rarely 0 - 20%, Sometimes 21-60%, Routinely 61-100%. 54% of physicians and 64% of HP_Yes indicated that patients routinely ask about exercise. However patients rarely ask HP_No about exercise (73%); a significant difference in distribution of proportions (χ^2^ = 31.48, p < .0005). Post-hoc pairwise comparisons using the z-test, with Bonferroni correction, showed no statistical differences between physicians and HP_Yes, but significant differences with HP_No in two categories: Rarely & Routinely. (B) Cumulative percent of responses when asked about familiarity with the ACSM exercise guidelines for pALS. Higher number indicate more familiarity: e.g. 1-2 indicate no to little familiarity, 6-7 moderate to strong familiarity. Therefore a shift of the curve rightward indicates greater familiarity. A Kruskal-Wallis H test revealed significant differences in medians (H(2) = 14.908, p = .001) which are indicated by arrow heads. Pairwise comparisons were performed using Dunn’s procedure with a Bonferroni correction for multiple comparisions. This analysis revealed statistically significant differences in median familiarity scores between HP_No (1.0) and Physician (2.5) p = .005, and HP_No and HP_Yes (4.0) p < .0005, but not between Physician and HP_Yes groups.

### Intentions toward exercise counsel

When participants were asked if they would provide exercise counsel to pALS in the future, their responses were divided (Figure 2A). Intentions were not the same for the three exercise modalities (χ^2^(2) = 33.977, p < .0005) with stronger intentions for flexibility compared to strength (p = .013) or aerobic (p = .027) training. There were no significant differences in distribution of intention scores for strength and aerobic training (p = 1.0). Across the three exercise modalities, 24% of respondents were unlikely (very unlikely + unlikely) while 45% were likely (very likely + likely) to provide exercise counsel. The unlikely intentions are almost exclusively from non-physician health professionals that indicated exercise counsel is not within their scope of practice (*HP_No_*, Figure 2B). Consequently, there were differences in intentions between the three groups (χ^2^(2) = 31.555, p < .0005) with lower mean ranks for intentions in the HP_No_ group compared to either Physicians (p < .0005) or *HP_Yes_* (p < .0005). Participants in the physician and *HP_Yes_* groups had similar fractions of respondents classified as ‘likely’ to provide exercise counsel (48% of physicians and 57% of HP_Yes_, χ^2^(1) = .382, p = .536) and average intentions were considered to have a positive valence (5.4 physicians, 5.8 *HP_Yes_*). The *HP_No_* group was not considered in the next phase of analysis given that they consider exercise counsel outside of their scope of practice and the concomitant low intention values.

**Figure 2.**
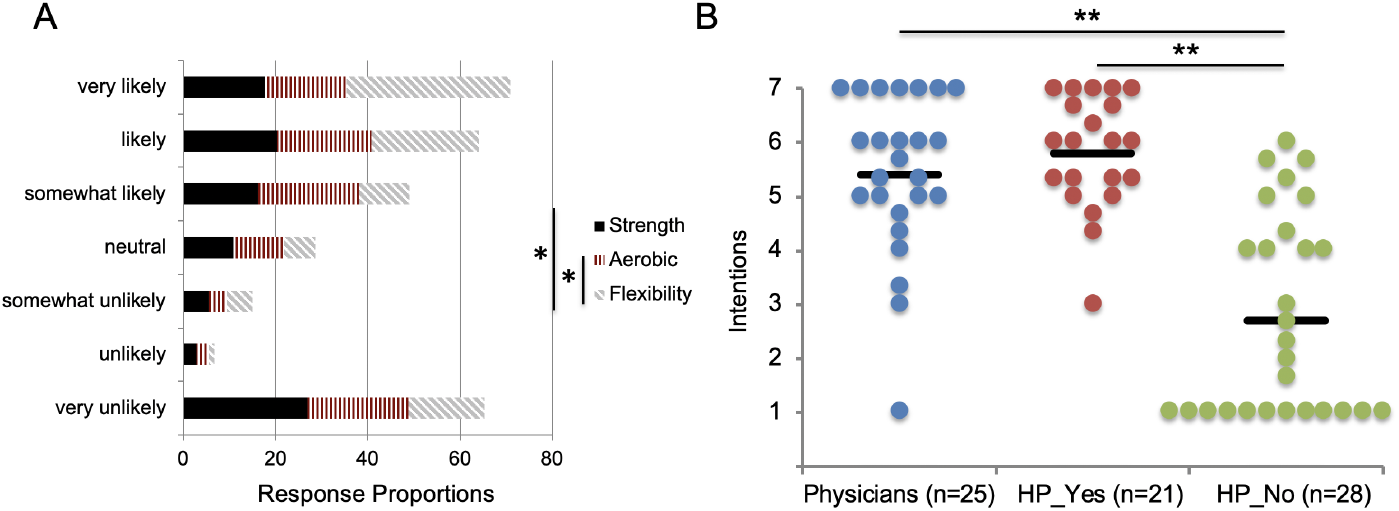
Intentions toward providing future exercise counsel. (A) Stacked intention responses for each of the three exercise modalities. The proportions are calculated within each exercise category, e.g. sum of black bars = 100%. On average about 24% of respondents had low (responses very unlikely-likely or 1-2) and 45% had high intentions (likely-very likely or 6-7). The distribution of responses is not the same for the 3 exercise modalities: Friedman two-way analysis of variance by ranks for repeated measures, N = 73 χ^2^(2) = 33.977, p < .0005. Pairwise comparisons using Wilcoxon signed-rank test * p < .05. (B) Average intention score across 3 exercise modalities divided into three respondent groups. Black bars show mean. Distributions of intention were not similar across groups, as assessed by visual inspection of the dot density plots, so difference in mean ranks was tested using a Kruskal-Wallis H test χ^2^(2) = 31.555, p < .0005. Dunn’s procedure for post-hoc pairwise comparisions revealed differences between HP_No (mean rank = 19.77) and both other groups (Physicians = 46.30, HP_Yes = 50.67), which were not different from each other. ** p < .0005.

### Theory of Planned Behaviour constructs and correlations with intentions

Attitudes towards exercise for people living with ALS were positive (i.e. >5.0), while subjective norms and perceived behavioral control were neutral-to-positive (Figure 3). Exercise modality had a statistically significant effect on the values of the three constructs of the TPB: Attitude F(2,90) = 22.863, p < .0005, partial *η*^2^ = .337; Subjective Norms F(2,82) = 17.297, p < .0005, partial *η*^2^ = .297; Perceived Behavioral Control F(2,90) = 22.546, p < .0005, partial *η*^2^ = .334. For all three constructs, the average values for flexibility were greater than those for the other two modalities, which were not different from one another (Bonferroni post hoc tests for pairwise comparisons, details in figure legend). There were no differences in responses between physicians and the *HP_Yes_* group. While attitudes toward exercise were generally positive in both groups, they were not correlated with intention to provide exercise counsel (Table 2). Intention was significantly correlated with subjective norms and perceived behavioral control. Multiple regression analyses were run to determine which of the TPB variables had predictive value for intentions. Initial models using the profession group (physicians or *HP_Yes_*) revealed that this variable did not have significant predictive value; therefore, analyses were done without separation into professional groups. The multiple regression models predicted intentions across the three exercise modalities with large effect sizes [29]: Strength F(3,39) = 13.189, p < .0005, adj. R2 = .47; Aerobic F(3,38) = 7.211, p = .001, adj. R2 = .31; Flexibility F(3,40) = 8.885, p < .0005, adj. R2 = .36. However, only subjective norms and/or perceived behavioral control had statistically significant regression coefficients as shown in Table 2. The proportion of variance in intentions accounted for by the regression models ranged from 31 – 47%, indicating variance in intentions may be accounted for by other factors.

**Figure 3.**
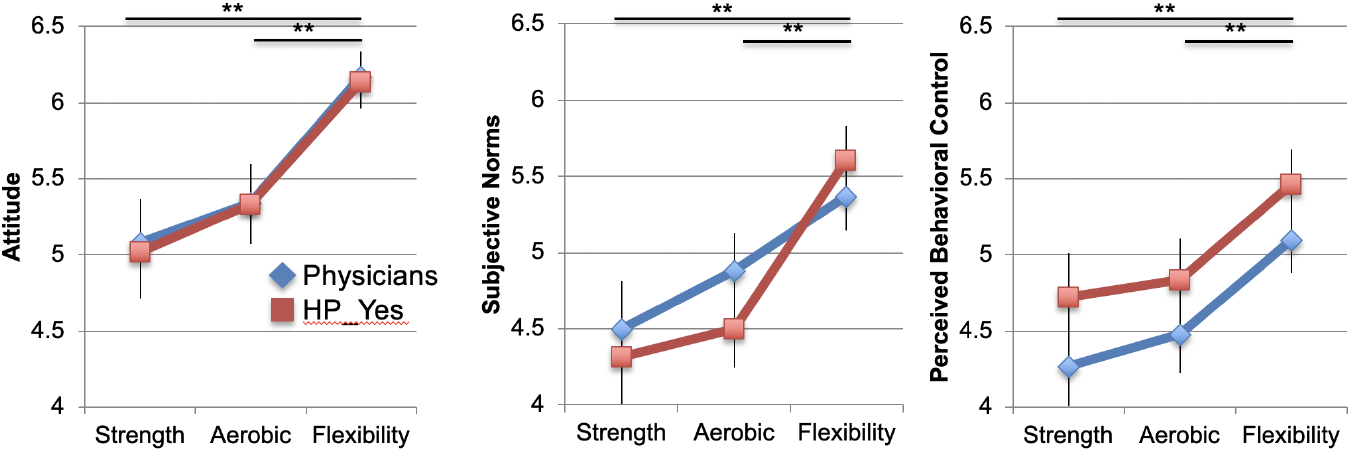
Exercise modality effect on TPB constructs. Two-way mixed ANOVAs showed no statistically significant two-way interactions for the three measures. There were no statistically significant main effects for the between subjects factor, i.e. physicians versus HP_Yes: Attitude F(1,45) = .013, p =.91; Subjective Norms F(1,41) = .120, p = .731; Perceived Behavioral Control F(1,45) = 1.397, p = .243, partial *η*^2^ = .03. There was a main effect for the within-subjects factor, exercise modality. Bonferroni post hoc tests were done for paired comparison of exercise modalities. Attitude: Flexibility vs Strength and/or Aerobic p < .0005, Strength vs Aerobic p = .182; Subjective Norms: Flexibility vs Strength and/or Aerobic p < .0005, Strength vs Aerobic p = .343; Perceived Behavioral Control: Flexibility vs Strength and/or Aerobic p < .0005, Strength vs Aerobic p = .364. Data points are marginal means with standard error.

**Table 2.** TPB correlations with intentions and multiple regression analyses. *β* = unstandardized regression coefficient. Significance at the .05(*) or .01 (**) levels.

|  | Attitude | SN | PBC |
| --- | --- | --- | --- |
| *Strength* |  |  |  |
| Physicians | .365 | .571** | .658** |
| HP_Yes | .454* | .500* | .601** |
| $\beta$ (± Std error) | .299 ± .214 | .471 ± .211* | .374 ± .190 |
| *Aerobic* |  |  |  |
| Physicians | .270 | .492* | .719** |
| HP_Yes | .206 | .661* | .293 |
| $\beta$ (± Std error) | -.027 ± .199 | .557 ± .185** | .371 ± .171* |
| *Flexibility* |  |  |  |
| Physicians | .223 | .495* | .505* |
| HP_Yes | .238 | .374 | .509* |
| $\beta$ (± Std error) | .079 ± .199 | .135 ± .165 | .649 ± .163** |

### Other measures correlated with intentions

There were differences between the physician and *HP_Yes_* groups in the non-TPB items that were associated with intentions (Table 3). Intentions of physicians to provide modality specific exercise counsel had strong positive correlations with the ability of their patients to perform exercises outlined in the ACSM guidelines. A significant correlation for this item was present only for the flexibility mode in the *HP_Yes_* group. Physician intentions for strength and aerobic exercise were also strongly correlated with their recall of the number of patients that were given exercise counsel over the previous month, *i.e*. previous behavior. Intentions of the *HP_Yes_* group to provide strength and aerobic exercise counsel had strong positive correlations with their familiarity with the ACSM guidelines. The proportion of pALS asking for exercise advice was not significantly correlated with intentions, except in the case of flexibility exercise in the physician group.

**Table 3.** Non-TPB correlations with intentions to provide exercise counsel. Nonparametric (Spearman) correlations, significance at the .05(*) or .01 (**) levels.

|  |  | Strength | Aerobic | Flexibility |
| --- | --- | --- | --- | --- |
| *Familiar with ACSM guidelines* |  |  |  |  |
|  | Physicians | .295 | .322 | .256 |
|  | HP_Yes | .594** | .599** | .296 |
| *Proportion of pALS that ask about exercise* |  |  |  |  |
|  | Physicians | -.011 | .153 | .555** |
|  | HP_Yes | .249 | .180 | .206 |
| *Proportion of pALS capable of exercise according to ACSM guidelines* |  |  |  |  |
|  | Physicians | .712** | .630** | .603** |
|  | HP_Yes | .119 | .258 | .518* |
| *Number of pALS advised about exercise in the last month* |  |  |  |  |
|  | Physicians | .573** | .490* | .264 |
|  | HP_Yes | .356 | .245 | .385 |
| *Team involvement in exercise prescription for pALS* |  |  |  |  |
|  | Physicians | -.048 | -.371 | -.192 |
|  | HP_Yes | .576** | .402 | .317 |

### Reasons for not providing exercise counsel

Five categories of responses were determined when respondents were given the opportunity to list three reasons for not providing exercise prescription to pALS (Table 4). The most frequent categories of response were patient centered (e.g. tolerance, adherence and interest) closely followed by responses that reflected uncertainty about whether exercise counsel was the respondent’s responsibility and whether they had sufficient expertise. Responses in these two categories accounted for 57% of the total number of responses. Examples of representative responses for the first category included, “Most patients need to save their energy to do the things they enjoy” and “Patient not interested”. Responses encoded in the second category included “I believe another team member has it covered” and “Not my expertise”. The category with the third most frequent responses included responses “Other priorities in consultation”, “Don’t have the necessary adapted equipment” and “Time”. The next most frequent response category included “There are not well accepted guidelines on what the exercise prescription in ALS should be” and “Not sure if strength exercises really help the patient”. The final category was used for responses such as “I have not covered ALS clinic in the last month” and “I am mostly retired”. The greatest differences when comparing the physicans and *HP_Yes_* groups were in the second and fourth ranked categories. A greater proportion of *HP_Yes_* respondents were uncertain about their expertise to provide exercise counsel or whether the role fell to another member of the multidisciplinary team. A greater proportion of physician responses centered on the issue of evidence for benefit of exercise.

**Table 4.** Open-ended responses for *not* providing exercise counsel. A total of 165 responses were classified into five categories. The percent of responses in each category are given along with raw numbers.

| Category | All Responses % (N) | Physicians | HP_Yes |
| --- | --- | --- | --- |
| 1 Patient tolerance, adherence, interest | 29 (48) | 31 (17) | 30 (31) |
| 2 Lack of confidence or competence, another team member's responsibility | 28 (46) | 13 (7) | 32 (33) |
| 3 Lack of time, space, resources, other priorities | 16 (27) | 22 (12) | 15 (15) |
| 4 Lack of evidence, no benefit of exercise | 15 (24) | 22 (12) | 11 (11) |
| 5 Other | 12 (20) | 13 (7) | 13 (13) |

## Discussion

It is clear from this study that the topic of exercise is routinely brought up by people living with ALS (pALS) in discussions with their health professionals, especially those who consider exercise prescription within their scope of practice. It is also clear that these health professionals have strong intentions to provide exercise counsel to pALS, in contrast to our proposal that intentions would be low to moderate. The exercise advice given by physicians is not likely to originate from the ACSM guidelines [1] because they were largely unknown and not associated with their intentions. However, the intentions of non-physician health professionals were strongly associated with their familiarity of the ACSM guidelines. Overall there is a clear preference towards flexibility exercise for intentions, and the proximal predictors: attitudes, subjective norms and perceived behavioral control. We found that potential barriers to providing exercise prescription are in the domains of subjective norms and perceived behavioral control, though open-ended responses from physicians also included lack of evidence.

At the time of data collection guidelines from the American Academy of Neurology (AAN) [30] and the European Federation of Neurological Societies (EFNS) [31] represented the best practise for ALS management. These guidelines for the care of pALS were largely silent on the utility of exercise as a disease modifying therapy. Both guidelines included a reference to Class III evidence for exercise as a treatment for pALS experiencing spasticity [32] but came to slightly different conclusions: insufficient evidence to recommend (AAN), and a consensus opinion that exercise is good clinical practice (EFNS). It is likely that our sample of health professionals from Canada were more familiar with the AAN guidelines, though we did not measure this. Another report available at the time was the update to the original 2008 Cochrane review “Therapeutic exercise for people with amyotrophic lateral sclerosis or motor neuron disease” [14]. Only two randomised control trials met the inclusion criteria for the review, both indicating that modest load resistance exercise had the potential to slow functional decline [32, 33]. The authors of the review point out that there was a complete lack of randomised or quasi-randomised clinical trials on the effects of aerobic exercise in pALS [14] and concluded with the familiar “*More research needed*”, an imperative also in the AAN Practise Parameter update [30]. Thus the background professional context at the time of the survey may be summarized as *cautiously optimistic* about the potential benefit of exercise and this is reflected in the positive attitudes towards exercise in our study.

As proposed we found a clear bias in favor of flexibility exercise. Health professionals reported that their clinics highly encouraged flexibility exercise for pALS (Supplementary Table 5) and the Theory of Planned Behavior (TPB) constructs reflected a clear bias in favor of flexibility over strength and aerobic exercise. In fact, in all measures strength training exercise was least favoured. What are the potential reasons for the bias in favor of flexibility over strength training? It could be the lingering perception that flexibility exercise does no harm, whereas strengthening could make things worse or exacerbate spasticity. Upper motor neuron signs in ALS diagnosis typically include spasticity and hyperreflexia [34] and the mainstay for non-drug management of spasticity and preventing contractures, has been stretching and flexibility exercise [22, 35]. The best evidence at the time was that individuals with neurological conditions, received no clinical benefit from stretching when assessing joint mobility and spasticity [36]. A recent update of that systematic review concluded that no further research is necessary since the evidence that stretch is ineffective, is of a high quality and additional studies are unlikely to change the findings [37]. However the systematic review did not include participants with ALS; though it is unlikely that pALS would respond differently to individuals with other neurological conditions like stroke, multiple sclerosis and spinal cord injury associated with upper motor neuron syndrome and spasticity. The absence of studies examining stretch interventions in participants with ALS was not because the search process excluded this population, but because there were no trials that met the inclusion criteria. There was a lack of randomized or quasi-randomized clinical trials examining the effectiveness of stretch for spasticity, contractures or range of motion in pALS. Despite this lack of evidence for the effectiveness of stretch in pALS, intentions and attitudes of Canadian health professionals to promote flexibility exercise are high. This discrepancy between evidence and intentions may reflect that clinical experience, suggests benefits to pALS other than reducing spasticity and maintaining range of motion. For example, health professionals may believe that stretch for pALS on a regular basis over the course of their lives has clinical and psychological benefits yet to be measured.

### Limitations

It is important to acknowledge that there was a delay between initial data collection and final analysis and write up. This means that current intentions may not be reflected in this cross-sectional survey and the sampling frame used may not accurately reflect the current constituents of the academic ALS clinics across Canada (see supplementary data at https://doi.org/10.6084/m9.figshare.8044622.v2). The convenience sample of voluntary respondents and an estimated response rate of 48%, is subject to selection and response bias. However, we received responses from a heterogeneous group of health professionals that reflects the multidisciplinary personnel working in the ALS clinics perhaps limiting the threat of response bias. Our division of non-physician health professionals into two groups: *HP_Yes_* and *HP_No_*, was based on self-report without asking about professional training that addressed exercise prescription for pALS. Another shortcoming of the study design was the absence of any measure of the health behavior of interest: exercise prescription or advice. Instead we chose to measure the TPB construct of intentions as a predictor of behavior, and this is not always reliable [38]. Finally, when pilot testing and finalizing the survey instrument we attempted to make measures brief to minimize time for responses yet assess the main TPB constructs. The low internal consistency for PBC items suggests the assessment of this construct can be improved.

### Implications and Future Directions

Canadian health professionals have clear intentions to provide exercise counsel for pALS. These intentions are moderated by the medical maxim, *Primum non nocere* (i.e. “Do no harm”) indicated by their primary reason given for not providing exercise counsel, concern for the patients. The results of the most recent clinical trial of exercise for pALS focussed on safety, tolerability and compliance and found that all exercise types were safe and well tolerated [16]. There was no evidence in this trial that the exercise slowed the rate of functional decline, but there was a trend toward a reduced fall risk with strength and aerobic exercise. Most importantly, there was no evidence of exacerbation of outcomes related to patient function that were attributed to exercise. A barrier for physicians at the time of data collection was lack of evidence of benefit of exercise. In addition to the results of [16], where no statistical benefit was shown, two other recent exercise trials have reported reduced rates of functional decline associated with aerobic or combined aerobic and strengthening exercise [15, 39]. Ideally a metaanalysis and update of the Cochrane review [14] will provide clarity on the issue of benefit of exercise for pALS. There is sufficient evidence that strength training is an important component of healthy aging [40] and the accumulated evidence for pALS indicates that at best, it slows rather than speeds the expected declines in function associated with ALS and is safe. In considering evidence for promoting strength training, ALS health professionals should be mindful of the [lack of] evidence for flexibility and it’s status as a widely accepted prescription for ALS management [22]. Finally, perhaps it is time for the multidisciplinary ALS clinics in Canada to discuss scope of practice for physical activity and exercise prescription so that efforts to build competence and confidence can be developed for the target health professionals.

## Conclusions

The current study provides empirical evidence that while intentions are high to provide exercise counsel for pALS, there is a positive bias towards flexibility that does not originate from high quality clinical trials. Intentions toward exercise prescription in the strengthening domain appear to be lower than current evidence suggests, especially when considered alongside promotion of flexibility. More high quality evidence has accumulated in the past four years and it is expected that future updates to Practice Parameters and clinical management guidelines will include explicit recommendations for health professionals responding to pALS who enquire about the role of physical activity and exercise after diagnosis.

## List of Abbreviations

pALS: people living with Amyotrophic Lateral Sclerosis
HP: all non-physician Health Professionals
*HP_Yes_*: non-physician Health Professionals who answered ‘Yes’ to indicate exercise prescription is within their scope of practise
*HP_No_*: non-physician Health Professionals who answered ‘No’ to indicate exercise prescription is outside their scope of practise
ACSM: American College of Sports Medicine
EIM: Exercise is Medicine^®^
RCT: Randomized Controlled Trial
TPB: Theory of Planned Behavior
SDT: Self-Determination Theory
SN: Subjective Norms
PBC: Perceived Behavioural Control
AAN: American Academy of Neurology
EFNS: European Federation of Neurological Societies

## Declarations

### Ethics approval and consent to participate

The procedures followed were approved by the University of Alberta’s Research Ethics Board (Pro00047048) and were in accordance with the ethical standards outlined in the “Tri-Council Policy Statement” and the Helsinki Declaration of 1975, as revised in 1983. Potential respondents were informed that survey completion was considered implied consent prior to participation.

### Consent for publication

Not applicable

### Availability of data and material

The survey instrument and data representing the final analysis are available on Figshare.com at https://doi.org/10.6084/m9.figshare.8044622.v2.

### Authors’ contributions

Initial conception and design (KEJ, TRB), revision of design and measurement tools (KEJ, TRB, VDBH), data analysis (KEJ, TRB), interpretation of data (KEJ, TRB, VDBH), drafting manuscript (KEJ) manuscript revision and final approval (KEJ, TRB, VDBH). The data were collected and separate analysis was done for a thesis submitted in partial fulfillment of the requirements for the degree of Master of Science by Aaliya Merali [41]. Repeated attempts were made to contact Aaliya and provide her the opportunity to read and revise the manuscript, but we were unsuccessful and therefore have not included her as a coauthor.

## Acknowledgements

Thank you to Dr Lorne Zinman for assistance in dissemination of the survey through the Canadian ALS Research Network. Thanks also to the administrative personnel at the provincial ALS clinics for assistance in enumerating the potential respondents and disseminating the survey advertisements and reminders. Thanks to Drs Eleanor Stein and Joanna Auger for constructive discussion and comments on the manuscript.

---

**Table 5.** *Supplementary* Perceived Behavioral Control: individual items

|  | Strength | Aerobic | Flexibility |
| --- | --- | --- | --- |
| *It is easy to provide exercise prescriptions for pALS.* |  |  |  |
| Physicians | 4.0±1.9 | 4.8±1.9 | 4.8±1.5 |
| HP_Yes | 4.4±1.7 | 4.3±1.6 | 5.6±1.3 |
| *I am confident in providing exercise counsel to pALS* |  |  |  |
| Physicians | 4.8±1.7 | 5.0±1.5 | 5.4±1.5 |
| HP_Yes | 4.9±1.6 | 5.2±1.6 | 5.5±1.9 |
| *My clinic accommodates exercise advise for pALS (i.e. has the facility, space, equipment)* |  |  |  |
| Physicians | 3.0±2.1 | 2.9±2.2 | 3.6±2.5 |
| HP_Yes | 3.8±2.1 | 4.0±2.0 | 4.3±2.1 |
| *My clinic encourages exercise counsel for pALS* |  |  |  |
| Physicians | 4.4±2.0 | 4.8±1.8 | 5.9±1.1 |
| HP_Yes | 4.9±1.8 | 5.2±1.7 | 5.9±1.5 |

## Additional Files

Additional file 1 — Sample additional file title

Additional file descriptions text (including details of how to view the file, if it is in a non-standard format or the file extension). This might refer to a multi-page table or a figure.

Additional file 2 — Sample additional file title Additional file descriptions text.

